# Detecting Amino Acid Variants Using Next-Generation Protein Sequencing (NGPS)

**DOI:** 10.1101/2024.12.17.629036

**Authors:** Mathivanan Chinnaraj, Jianan Lin, Kristin Blacklock, Eric Hermes, Michael Meyer, Douglas Pike, John Vieceli, Ilya Chorny

**Author notes:** Equally contributing first authors (Mathivanan Chinnaraj, Jianan Lin). Remaining authors in alphabetical order.

## Abstract

Next-Generation Protein Sequencing^™^ (NGPS^™^) is a single-molecule approach for characterizing protein variants, offering detailed insight into proteoforms and amino acid substitutions not easily discerned by mass spectrometry. The novel data type produced by NGPS, which is based on binding of N-terminal amino acids by fluorescently tagged recognizer proteins, requires the development of new data analysis methods and bioinformatic tools. Here, we present ProteoVue^™^, a comprehensive bioinformatics pipeline for Single Amino Acid Variant (SAAV) detection and quantification using the Quantum-Si Platinum^®^ NGPS platform. ProteoVue integrates multiple analytical components, including robust pulse-calling, recognition segment detection, fluorescence dye classification, and a neural network-driven kinetic signature database for pulse duration prediction. These components feed into a scoring-based alignment and clustering framework that enables accurate variant calling within binary peptide mixtures. We demonstrate that ProteoVue recovers expected variant ratios across diverse substitution types including residues that lack direct amino acid recognizers. While some extreme cases remain challenging, the pipeline consistently captures the key kinetic features required for variant discrimination, underscoring its potential as a versatile and powerful tool for proteomic studies. As NGPS technology matures and recognizer libraries expand, ProteoVue provides a foundation for increasingly refined variant analysis in basic research, biomarker discovery, and clinical applications.

## Introduction

Quantum-Si’s Next Generation Protein Sequencing (NGPS) [1] technology, which is commercially available via the Platinum platform, represents a significant advancement in proteomics. Like mass spectrometry (MS), NGPS offers a hypothesis-free measurement of the proteome. However, unlike ensemble protein analysis methods, NGPS enables real-time, single-molecule measurements of individual peptides [1]. NGPS can also measure features that are challenging for MS, such as isobaric amino acids and highly similar proteoforms [2]. Thus, NGPS is an orthogonal tool that is complementary to many established proteomics methods.

Proteoforms are protein variants arising from multiple sources of genomic, transcriptomic, and post-translational variation, including alternative splicing and post-translational modifications [3]. Due to the functional impacts of these variations, proteoforms play crucial roles in biological and disease mechanisms [3]. Individual proteomics techniques can struggle to capture the full diversity and complexity of proteoforms; therefore, multiple orthogonal tools may often be required [4]. NGPS uses single-molecule measurements to directly analyze individual protein molecules, providing detailed information on their modifications and variation. This capability opens new avenues for studying proteoforms.

The unique features of NGPS technology require the development of new data analysis methods and bioinformatic tools. Specifically, the use of N-terminal amino acid (NAA) recognizers to detect individual amino acids in a peptide (see Figure 1) requires specialized analysis methods that consider the unique biophysical properties of recognizer-based sequencing.

**Fig. 1:**
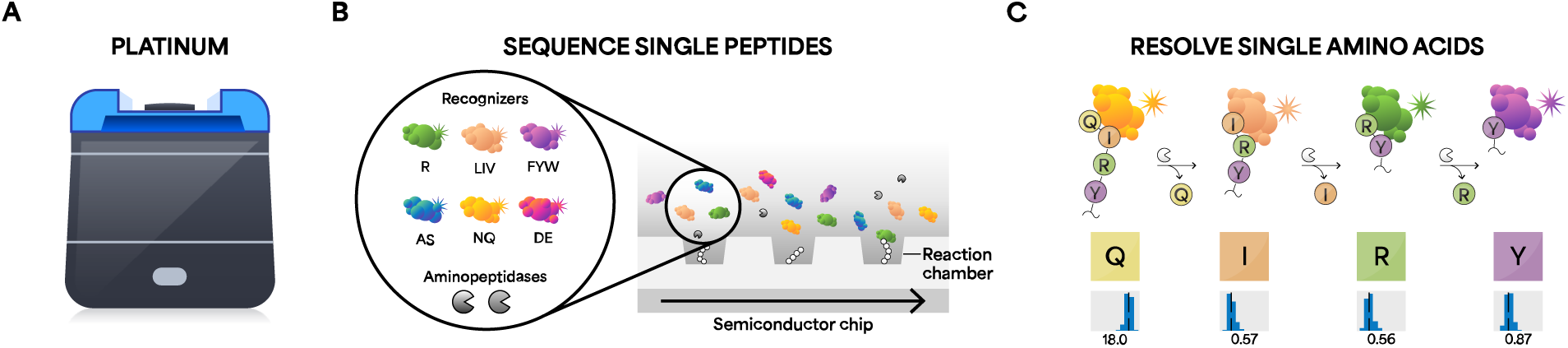
(A) The Platinum instrument sequences single peptide molecules with single amino acid resolution. (B) Sequencing kits include semiconductor chips, aminopeptidases, and six dye-labeled NAA recognizers that reversibly bind 13 target NAAs. (C) Binding of dye-labeled NAA recognizers generates kinetic information indicating which amino acid is being detected.

In this study, we demonstrate the detection of Single Amino Acid Variants (SAAVs) using NGPS. We showcase SAAV detection in binary mixtures of synthetic peptides and accurately predict the mixture ratio within a factor of ten of the expected value. The methods described herein also serve as a foundation for future studies focused on the bioinformatic analysis of proteomic variation in NGPS datasets.

## Materials and Methods

### Peptide Design

Synthetic peptides were designed for sequencing on Quantum-Si’s Platinum NGPS instrument. Each peptide followed the sequence RFNELXFDISRYLANK(N3), where X was substituted with F, W, R, M, N, A, or C, resulting in seven distinct sequences. Figure 2 is a schematic of the peptide design. A key requirement was that at least four amino acids N-terminal to the substitution could be recognized by the Quantum-Si V3 Sequencing Kit, with three of these four amino acids being uniquely identifiable. The choice of the sixth-position substitution was intended to capture kinetic variations associated with up to five preceding residues, arising from interactions between NAA recognizers and their downstream positions, facilitating variant calling in binary mixtures. Additionally, a C-terminal azido-lysine was incorporated to ensure compatibility with Quantum-Si’s Library Preparation Kit–LysC, enabling immobilization of the peptides on the semiconductor chip surface.

**Fig. 2:**
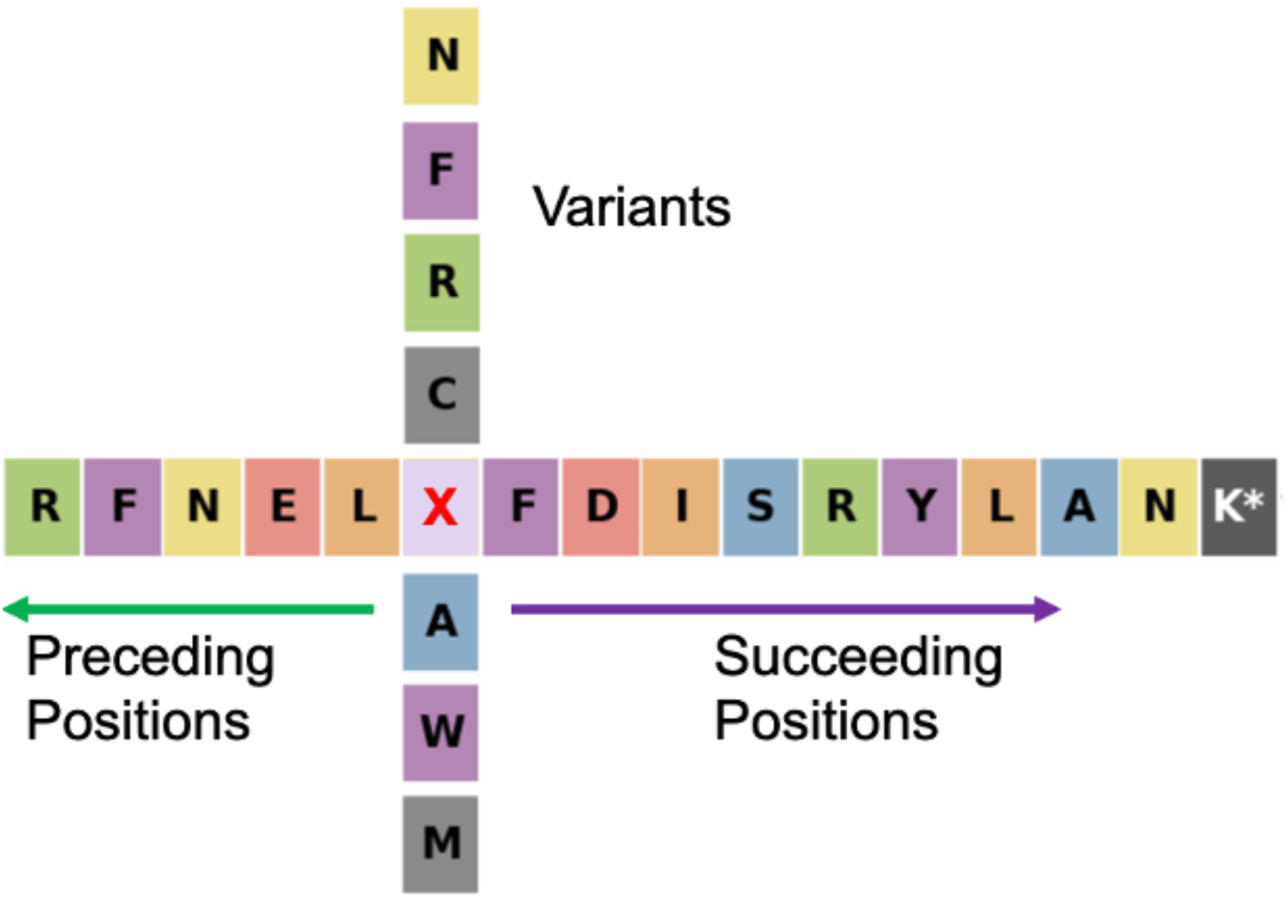
Design of peptide sequence with all 7 variants at 6^th^ position

### Peptide Synthesis

Seven peptides were synthesized by InnoPep (San Diego, CA), each supplied at 3 mg and with a purity greater than 95%. All peptides featured an N-terminal H, C-terminal carboxylic acid block NH2, as well as a C-terminal azido-lysine modification. They were initially reconstituted in DMSO to a concentration of 10 mM and stored at –20°C until linker conjugation. The azido-lysine modified peptides were then conjugated to the linker molecule from Quantum-Si’s Library Preparation Kit-LysC through strain-promoted alkyne-azide cycloaddition (SPAAC) click chemistry [5]. The peptides were then diluted to a final concentration of 20 µM in 100 mM HEPES, pH 8.0 (20% acetonitrile), and incubated with 250 µM Cetyltrimethylammonium bromide (CTAB) and 2 µM linker for 16 hours at 37°C. The conjugated peptide libraries were stored at –20°C until sequencing.

#### NGPS

Peptide sequencing was carried out on a Quantum-Si Platinum instrument. Approximately 200 pM of the conjugated peptides were loaded, followed by the removal of excess, unbound peptides. Sequencing utilized the V3 Sequencing Kit [6], which includes N-terminal amino acid (NAA) recognizers for 13 of the 20 canonical amino acids. Specifically, the V3 kit contains a set of six NAA recognizers for LIV, FYW, and R, as previously described [1], along with additional recognizers for AS, DE, and NQ. Each recognizer was uniquely labeled with a fluorescent dye, as described previously [1]. The binding and dissociation of these NAA recognizers to the immobilized peptides were monitored in real time as individual on-off events.

The V3 Sequencing Kit contains a combination of two aminopeptidases which cleave all twenty N-terminal amino acids at a controlled rate. The XP motif is the only exception and is not cleaved with the current V3 Sequencing Kit. NAAs from immobilized peptides are sequentially cleaved by aminopeptidases allowing the next amino acid to be exposed for NAA recognizers to bind. This process is repeated throughout the 10-hour run time.

Initially, seven sequencing runs were performed, each with one of the seven synthetic peptides. Additionally, binary mixtures of peptides were generated, asparagine (N) to alanine (A), (N6A), phenylalanine (F) to tryptophan (W), (F6W), arginine (R) to methionine (M), (R6M), and cysteine (C) to methionine (M), (C6M).

### Data Collection

Raw sequencing data was analyzed in real-time [1] on Platinum, producing pulse calls as output. The pulse calls were transferred to the cloud-based Platinum Analysis Software and analyzed using the Primary Analysis v2.8.0 (recognition segment detection), Peptide Alignment v2.8.0 (protein reference alignment), and ProteoVue v1.0.0 (two component Single Amino Acid Variant (SAAV) calling) workflows available in the Platinum Analysis Software (see Figure 3).

**Fig. 3:**
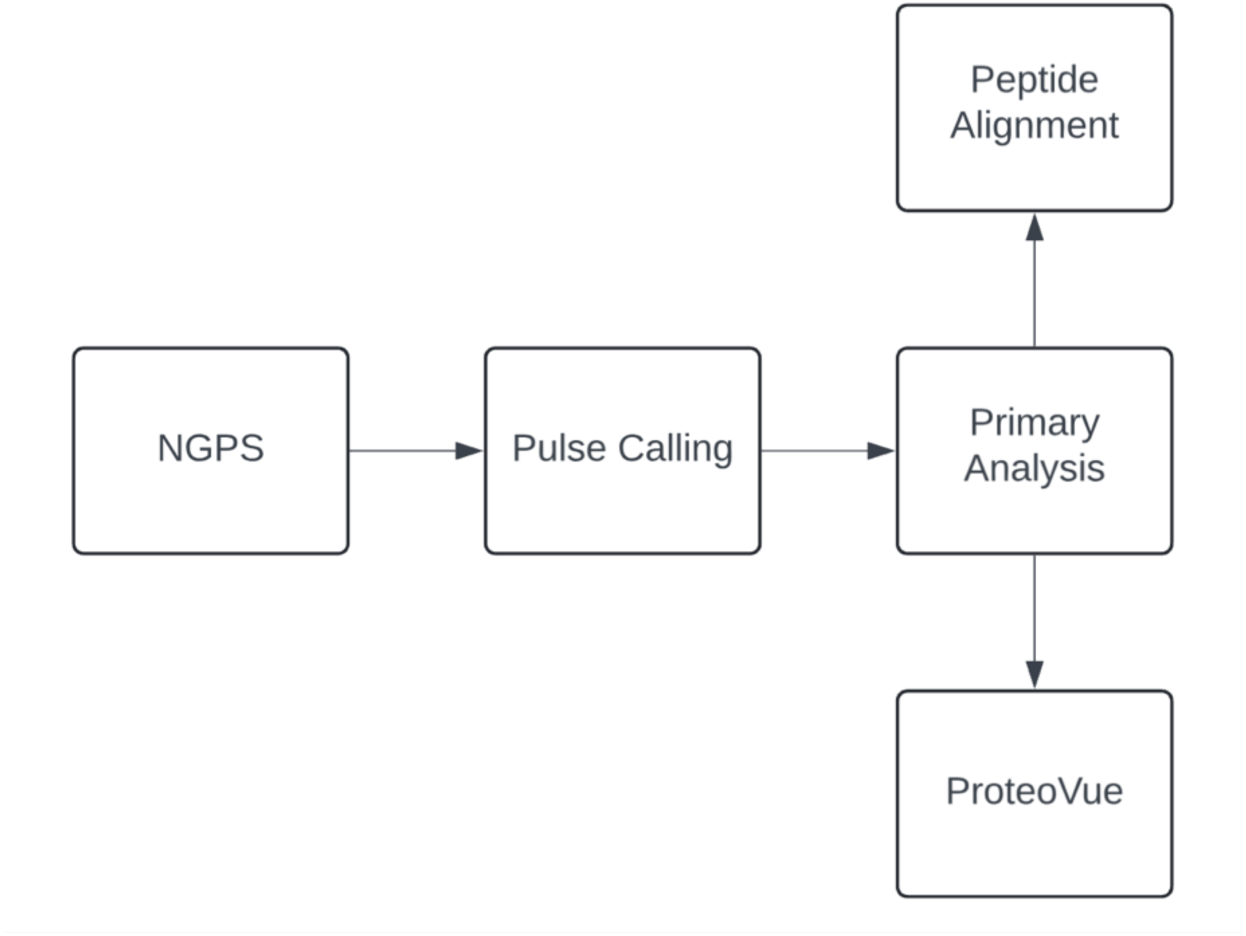
High level flow chart of analysis pipelines utilized in this study.

### Pulse Calling

The first stage of analysis is pulse calling, which occurs on the Platinum instrument. Here, raw signal data is analyzed to produce a set of pulse calls that can be fed into downstream signal processing and bioinformatics software.

Pulses are created when a recognizer binds to a peptide and its attached fluorophore is brought within range of incident excitation in the reaction chamber, resulting in an elevated fluorescent signal, which is recorded by the sensor [1]. The pulse caller is designed to detect these signals produced by binding events, and distinguish them from background noise, such as the fluorescence from freely diffusing recognizers also in the reaction chamber.

The pulse-calling algorithm operates on each reaction chamber independently and begins by estimating the statistical properties of the background noise. Once an estimate within acceptable error bounds is established, the algorithm processes new data frames in real-time. At each time point, it tracks whether the signal is attributed solely to the background noise or if it includes a pulse from a recognizer-NAA interaction. A transition from background to pulse occurs when an edge detection test identifies a significant signal shift compared to the background noise’s statistical distribution. Similarly, the transition from pulse back to background happens when recent frames of the signal match the background distribution. The algorithm continuously updates its background noise model in real time as new 60 ms frames are observed.

To account for the fact that detected pulses can represent either true recognizer-to-peptide interactions or transient noise spikes, a downstream filtering layer evaluates the significance of pulses. This evaluation considers factors such as pulse duration, intensity, and noise patterns across the duration of the run and the entire reaction chamber dataset.

Each pulse identified by the pulse caller represents a transient interaction between a recognizer and a peptide NAA. The pulse duration (PD) is governed by the dissociation kinetics of the recognizer-peptide complex. The inter-pulse duration (IPD) is the time from the end of one pulse to the beginning of the next pulse. Figure 4 is an illustration showing examples of PD and IPD. If the same recognizer-peptide interaction occurs in successive pulses, the IPD is governed by the association kinetics of the recognizer-peptide complex. Both PD and IPD can be modeled using exponential distributions derived from theoretical first-order reaction kinetics.

**Fig. 4:**
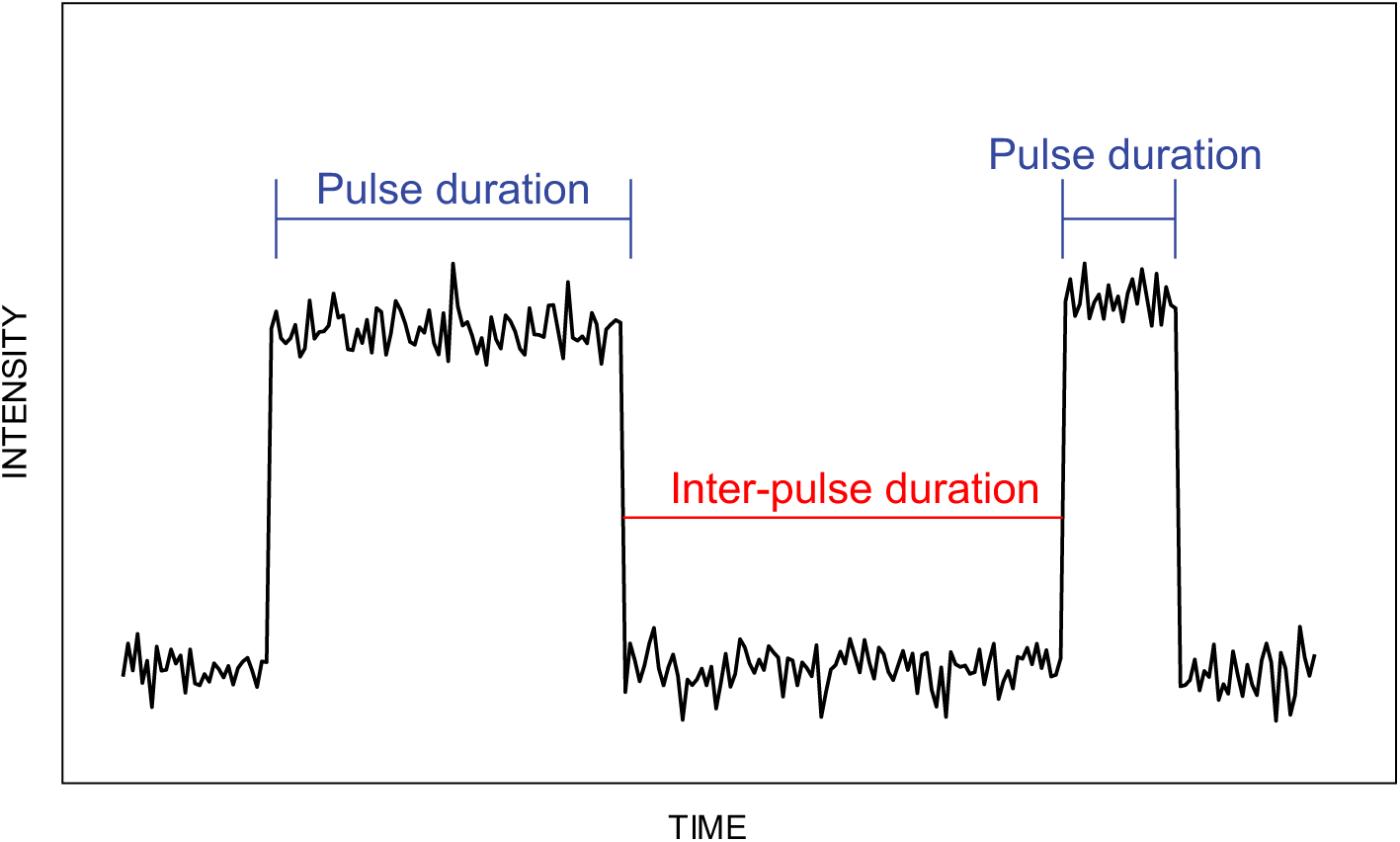
An illustration of two pulses, with the pulse durations (PD) highlighted in blue and the inter-pulse duration (IPD) between them highlighted in red.

### Primary Analysis - Trace Segmentation

As the sequencing reaction progresses, aminopeptidases progressively cleave N-terminal residues of the peptides in each reaction chamber, resulting in newly exposed N-terminal binding motifs whose recognizer affinities and kinetics largely differ from the previously exposed peptide state. The time to cleave an NAA range from approximately 10-40 minutes. Continuous pulsing associated with a single recognizer-peptide interaction is referred to as a recognition segment (RS). The goal of the trace segmentation algorithm is to identify these temporal segments by detecting where pulsing patterns change over the duration of the run.

The first step of trace segmentation is to identify boundaries where regions of active pulsing terminate, (i.e. where the peptide NAA state transitions from a residue that is detectable to one that is not). To do this, we scan a 30-pulse sliding window across the trace identifying significantly large gaps in time between any two pulses. The gaps are assessed by comparing the mean IPD of the 30 pulses in the window with the gap between the 30^th^ and 31^st^ pulse. If the gap is greater than 12 times the mean IPD of the preceding pulsing region, a split is made. The rationale for this rule is that given an exponential model of IPDs in the preceding region, 99.9999% of IPDs should be less than 12 times the mean, thus any gap greater than 12 times the mean IPD is very likely to indicate the end of the recognizer-NAA binding process of the previous 30 pulses. This procedure is then repeated by additionally scanning the trace in the reverse direction. By splitting the trace at these points, we divide the trace into “proto-RSs”, regions of active pulsing not containing any large gaps.

The algorithm next subdivides the proto-RSs into RSs, where each RS contains pulsing from only a single recognizer-peptide interaction. To accomplish this, we scan a sliding 60-pulse window over each proto-RS to identify potential pulse indices at which to split the proto-RS where the properties of the first 30 pulses differ significantly from the last 30 pulses, thus indicating a change in recognizer-NAA interaction after the 30^th^ pulse. The pulse properties assessed include those related to the kinetics of the interaction: pulse duration (PD) and interpulse duration (IPD), and those related to the fluorescent properties of the dye attached to each recognizer: dye intensity and fluorescence lifetime.

We compute p-values for four, independent 2-sample Kolmogorov-Smirnov (KS) tests [7], one for each pulse property, with the null hypothesis that the pulses in the first half of each window have the same distribution of the given property as pulses in the second half. The minimum of the four p-values is recorded for each potential pulse index split point across the proto-RS. If the minimum p-value across all potential split points is less than 10^-4^, the proto-RS is split into two proto-RSs at that pulse index. The p-value computations and splitting process is repeated on the resulting two proto-RSs and on any proto-RSs emanating from splitting those RSs, until no p-value meets the required threshold for further splitting. The resulting set of segments are called RSs.

### Primary Analysis - Dye Calling

As described in our previous publication [1], each recognizer is conjugated with a distinct dye that has a unique intensity and fluorescence decay lifetime. Platinum measures the fluorescence decay lifetime by sampling in two collection windows with different delay following laser illumination. In practice, the ratio of these two measurements (the "bin ratio") is used as a proxy for the fluorescence decay lifetime in downstream data analysis. The dye associated with each pulse is determined by a Gaussian Mixture Model (GMM) [8] classifier on the bin ratio and log intensity that is trained on-the-fly using many pulses sampled from across the chip and throughout the run. This GMM classifier is illustrated in Figure 5. After the GMM is fit, each cluster is associated with one of the dyes in the kit based on pre-calculated cross-run averages of the measured intensities and bin ratios for each dye.

**Fig. 5:**
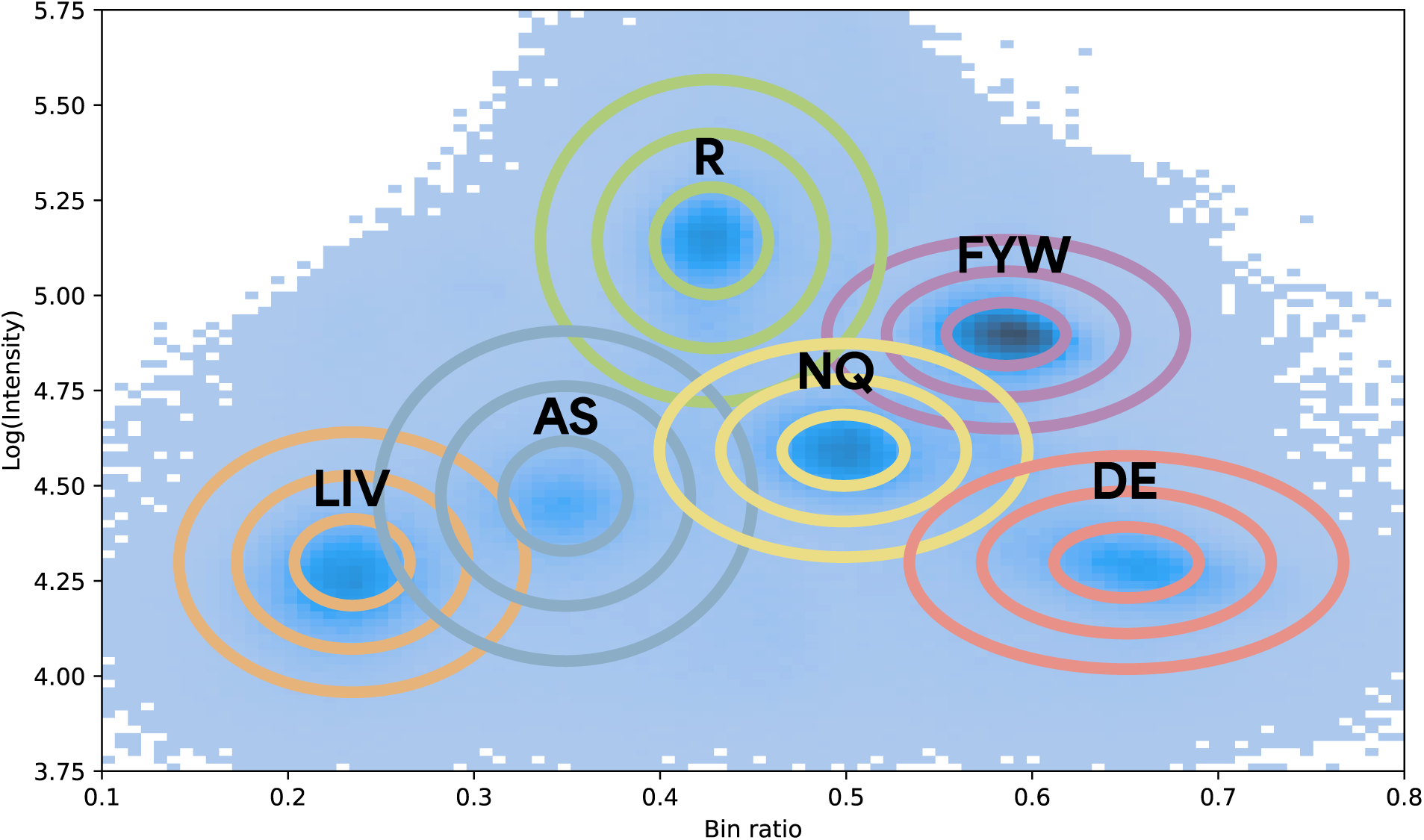
An illustration of the distribution of pulse bin ratios and log-intensities from a representative run, with the dye classification GMM overlaid on top represented as nested ellipses. Each ellipse represents one standard deviation of the normal distribution associated with each dye.

To determine which recognizer is associated with each RS, we use the GMM classifier to measure how consistent the distribution of pulse intensities and bin ratios within the RS are with each of the dyes in the GMM. This is referred to as the "dye purity", which is calculated using the following expression:

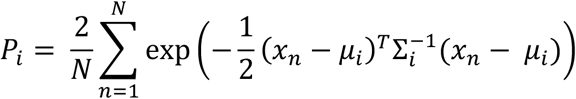

Where *P_i_* is the purity of the *i*th dye, *μ_i_* is the center of the *i*th dye’s normal distribution in the GMM, *Σ_i_* is the covariance matrix of the *i*th dye’s normal distribution, and *x_n_* is the position in log-intensity/ bin ratio space for the *n*th pulse in the RS. Purity values close to 1 indicate a high level of agreement between what is predicted for a particular dye and what is measured within the RS. The distribution of pulse bin ratios and intensities for each dye generally overlap somewhat. Consequently, even nominally “pure” RSs consisting only of pulses for a single dye will generally have non-zero purity values for all dyes. Using the GMM, it’s possible to predict what the observed purity should be for all other dyes:

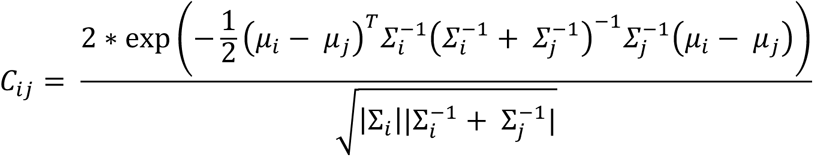

*C_ij_* is the purity of dye *j* we would expect to observe for a hypothetical RS consisting only of pulses whose bin ratio and intensity are sampled from the GMM distribution associated with dye *i*. To determine which recognizer to assign to each RS, we calculate and normalize the purity vector *P* of the RS and each column of *C* from the GMM, then calculate the distance between the *P* vector and each column of *C*. This distance is referred to as the dye composition distance. Each RS is labeled with the recognizer that has the shortest dye composition distance.

### Peptide Alignment

Once each RS has been labeled with a recognizer, each read is aligned against a panel of reference peptide sequences, with the highest-scoring alignment determining the inferred peptide identity of the read. The alignment algorithm matches each RS in a read to an amino acid residue in the reference such that the residue’s expected recognizer matches the recognizer label of the RS (Fig. 6). The score is based on the extent to which the observed pulse duration of the RS matches what is predicted by the kinetics database (see below) and the presence or absence of gaps between RSs that correspond to invisible states (states for which there is no recognizer) in the reference.

**Fig. 6:**
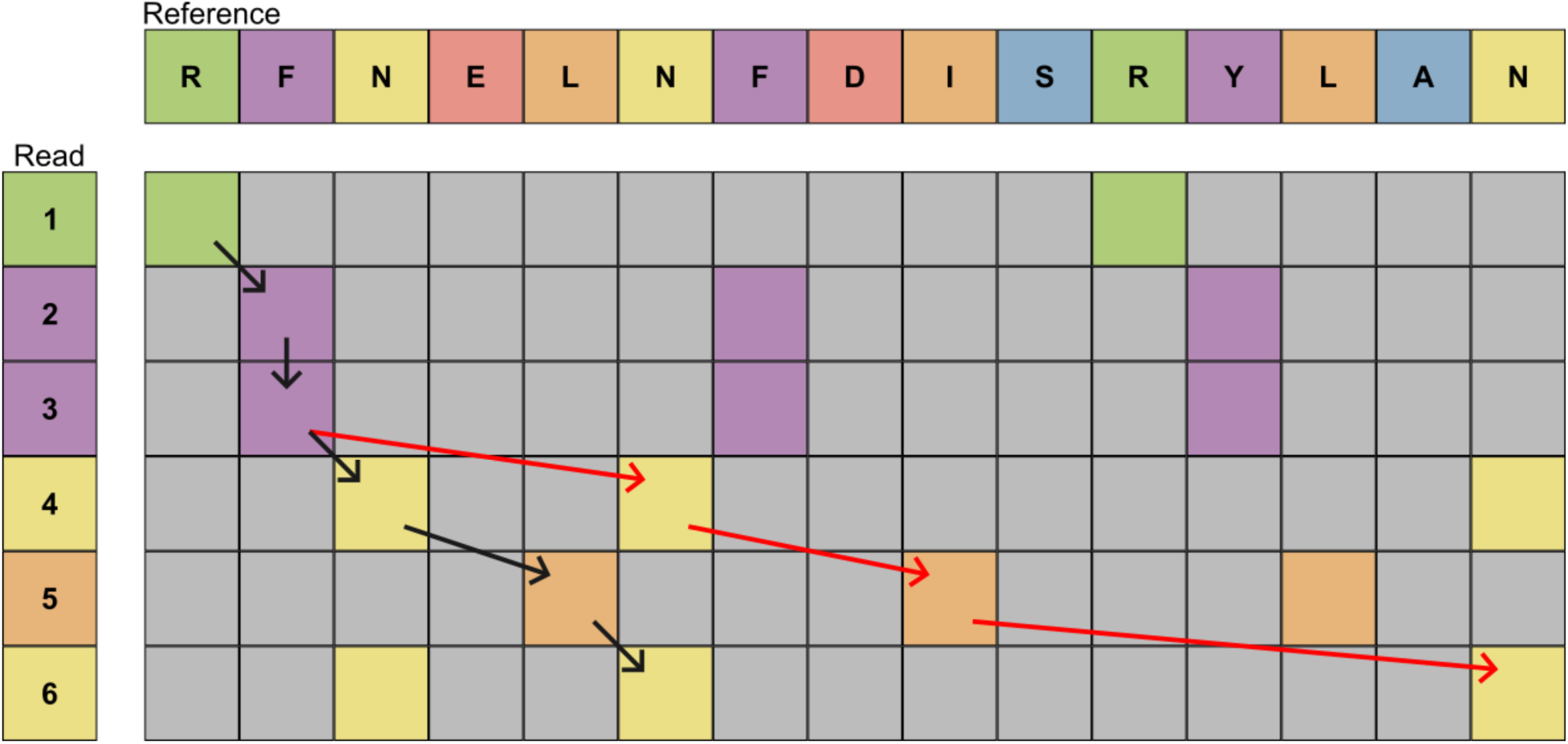
Illustrates an example alignment between a read and a reference peptide sequence. The color of each square indicates the presence of a recognizer match between the RS and reference state, with grey indicating that the recognizers do not match. The most likely trajectory is shown in black, with a second valid trajectory involving many deletions shown in red. States that do not have a matching recognizer are skipped by the aligner. In this alignment, RSs 2 and 3 are inferred to both correspond to the first F in the reference, which can occur when an RS is over-split. Additionally, none of the RSs in the read have a recognizer compatible with the E in the reference, which means the E state must be deleted in any valid alignment trajectory. For an alignment trajectory to be valid, it must align all RSs, but it does not need to reach the end of the reference sequence.

For a given pair of RS and visible reference state, the match score s_match_ is calculated as:

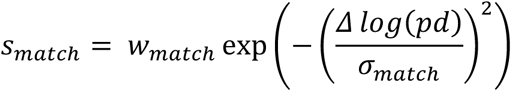

Where Δlog(*pd*) is the difference between the log observed pulse duration and the log expected pulse duration, σ_match_ is a scaling parameter for the penalty, and w_match_ is the match score weight. In our current implementation, w_match_ and σ_match_ are both chosen to be 1.

The aligner permits deletions, with states that have higher expected pulse durations receiving a larger deletion penalty. The deletion score is designed this way because we expect to miss some very short PD states due to the finite sampling rate of the instrument. The deletion score s_deletion_ for an ostensibly visible state that does not align to any RS in the read is given by

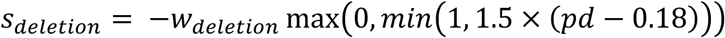

Where *pd* is the expected pulse duration in seconds of the state and w_deletion_ is the deletion score weight. In our current implementation, w_deletion_ is chosen to be 1. Because the number of amino acids sequenced will vary read-to-read, we do not apply a deletion penalty for any visible state following the one aligned to the final RS in the read.

Insertions are allowed by the aligner only if the recognizer of the inserted RS matches that of the RS immediately preceding or following it in the read. The aligner permits multiple adjacent RSs to align to the same state of the reference to ensure that an RS that was erroneously split into multiple RSs can still align. Additionally, RSs that are determined to be low quality are removed from the read prior to alignment. RSs with mean IPD above 42 seconds, dye purity below 0.15, or dye composition distance above 0.47 are considered low quality. This is meant to ensure that reads with misidentified, low-quality RSs can still align. If a valid RS is accidentally filtered out by this mechanism, the read may still align, albeit with a deletion penalty for the reference state that would have aligned to the removed RS.

The final contribution to the alignment score is referred to as the “gap score”, s_gap_, which rewards alignments in which the regions of no pulsing between RSs line up with states of the reference that we expect to be invisible, due to the lack of a specific recognizer for that state. Note that this is distinct from the gap penalty of DNA aligners, which refers to the penalty of contiguous insertions or deletions in the alignment that create gaps in aligned segments.

In our application, the gap score depends on the context of the alignment and needs to be bounded by realistic time frames that correspond to unrecognized amino acids. If two adjacent RSs align to the same reference state and there is a gap of at least 1200 seconds between them, then s_gap_ = -*w*_gap_ * 0.5, where *w*_gap_ is the gap score weight. If the two RSs are aligned to adjacent states and there is a gap of less than 300 seconds, then s_gap_ = *w*_gap_ * 0.5. If there is exactly one skipped residue in the reference (whether an invisible residue or a deleted visible residue) between two adjacent RSs and there is a gap of between 60 and 3000 seconds, then s_gap_ = *w*_gap_ * 0.5. Finally, if there are 2 or more skipped residues in the reference between adjacent RSs and there is a gap of greater than 1200 seconds, then s_gap_ = *w*_gap_ * 0.5. In all other cases, s_gap_ = 0. This formulation has the effect of penalizing alignments that imply a single state has been over split when the RSs aren’t adjacent in the read and rewarding alignments where the gap between RSs is consistent with skipped states in the reference, as these skipped states should create periods of no pulsing in the read.

One potential issue with the scoring algorithm described so far is that if a state is over split, this will increase the total number of RSs in the read, thereby increasing the maximum possible alignment score. Since downstream applications filter alignments to those above some minimum alignment score threshold, this scoring artifact has the effect of increasing the prevalence of over-split RSs, which introduces bias. To ameliorate this issue, each RS is given a score weight w_RS_ equal to 1/n, where n is the number of adjacent RSs in the read that have been labeled as the same recognizer. For example, if the RS recognizer sequence is ABBCB, then the state weights are 1, ½, ½, 1, 1. The final alignment score s_alignment_ is then the sum of each state’s match and gap score multiplied by the RS weight, plus the deletion penalty of any skipped state from the reference. For the purposes of weighting, each contribution to the gap score is associated with the subsequent RS in the read.

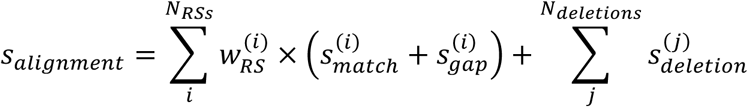

For a read to be a candidate for alignment, it must meet certain criteria that qualify it as a “high quality” read. For a read to be high-quality, at least 3 recognizers must be present among those assigned to the RSs. Additionally, the length of the read must be at least 4 after collapsing adjacent ROIs that share a recognizer (equivalently, the sum of RS weights must be greater than or equal to 4). These checks ensure that only reads containing enough information to reliably distinguish between peptide references are passed to the aligner.

### Kinetic Database

As described above, the alignment score relies on an expected PD to calculate the match score component *s_match_*. A PD prediction model was developed based on the hypothesis that PD is influenced by the sequence of the N-terminal residue and the subsequent four downstream residues, in the direction from N-terminus towards C-terminus. The trained model was then applied to predict pulse durations for all possible pentameric sequences starting with any of the 13 N-terminally recognized amino acids and containing any of the 20 canonical amino acids in each of the remaining four positions. This approach produced predictions for 2,080,000 (13 · 20^9^) unique sequences. In cases where a pulse duration has been previously measured for a corresponding pentameric sequence motif, the empirical value was used. In all other cases, the predicted value was used. Combining these results, a comprehensive kinetic database was generated, pairing each pentameric sequence with its corresponding pulse duration for use in scoring alignments against any pentameric sequence motif (7. 6C).

### Neural Network Model to Predict PDs

We developed a custom deep neural network model that integrates one-hot encoding of amino acid identities as well as sinusoidal positional encodings to incorporate sequence position into the feature representations without reliance on recurrent or convolutional operations [9]. The positional encoding function was adapted from the original Transformer model, using sine and cosine functions at different frequencies to produce a deterministic, continuous set of embeddings tied to each position index (Fig. 7B). Specifically, for an input sequence of length *5* and a feature dimension 𝐷𝑚𝑜𝑑𝑒𝑙, the encoding was defined elementwise as:

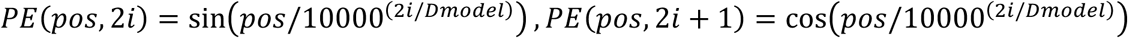

where 𝑝𝑜𝑠 is the position within the sequence and 𝑖 indexes the dimension. This positional encoding echoes a largely identity-independent effect that peptide sequence *position* has on pulse duration, which arises from the local environment surrounding each recognizer-bound peptide residue. As this per-position environment remains consistent from peptide to peptide, positional encoding imbues the model with a trainable parameter aimed at recapitulating the contribution of individual peptide positions on pulse duration that are inherent to the recognizer binding mode (Fig. 7B).

After applying the positional encodings to the input tensor, we concatenated these encoded amino acid features from the pentameric sequence motif with their corresponding pulse duration measurements. We then passed this combined feature set through a series of fully connected layers with gradually reduced dimensionality (128 → 64 → 16) to enhance computational efficiency and parameter regularization. Each of the first two layers was normalized using batch normalization [10] and regularized with a dropout rate of 0.4 [11]. ReLU activation functions were employed in all hidden layers to improve training stability and accelerate convergence [12]. Finally, a single-value regression estimate was produced by the output layer. By integrating sinusoidal positional encodings, dimensionality reduction, batch normalization, and dropout, the model aimed to achieve a more robust representation of sequence position while maintaining a compact and efficient architecture (Fig. 7C).

**Fig. 7:**
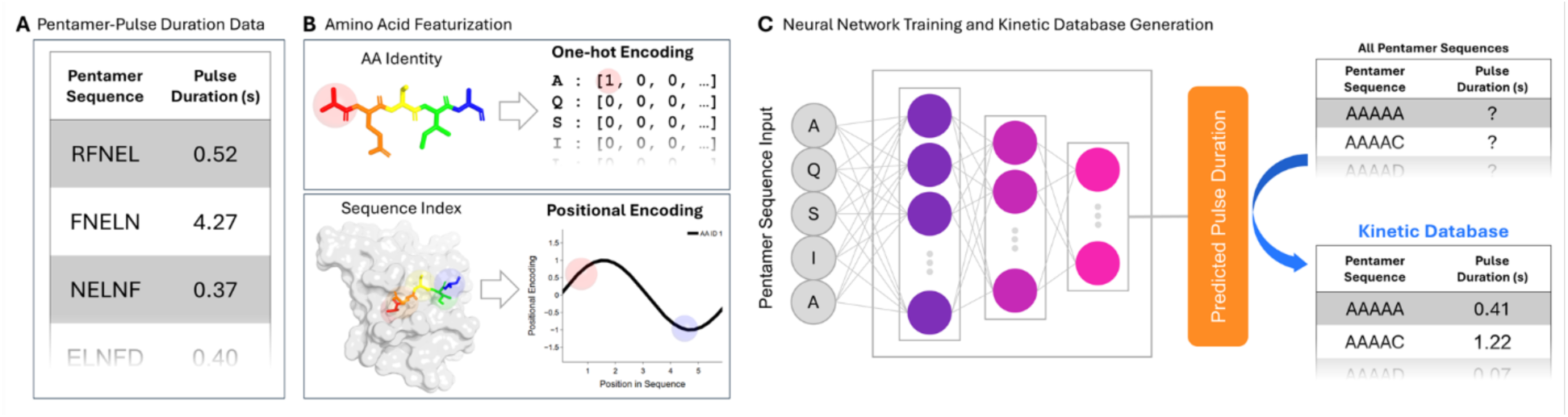
Architectural overview of the neural network used to generate the kinetic database. (A) We use pentamer sequence motifs and pulse duration data as inputs for training, validation, and testing. (B) The sequence-based input and featurization captures the identities and indices of each pentamer residue. Identities are incorporated using one-hot encoding, while sequence indices are encoded sinusoidally, ensuring each amino acid at each position has a unique representation within the positional encoding matrix. The importance of this positional context is founded in the recognizer:peptide (grey surface and rainbow sticks) binding mode, as one would expect an amino acid at the first position in the peptide (red sticks and spheres) to contribute more significantly to the overall pulse duration as compared to the same amino acid identity at position five (blue sticks and spheres) due to stronger intermolecular contacts. (C) We then train a fully-connected neural network model that is used to predict pulse durations for all possible pentamer motifs, thus generating a kinetic database.

### Training Data Curation

To generate the training data for the kinetic database neural network model, a series of runs comprising 30 proteins and 32 synthetic peptides were sequenced and aligned against their respective reference sequences using an iterative alignment approach that avoided circular dependencies. Entries were only included if a minimum of 100 recognition events per run were observed, constituting at least 10% of the alignment coverage at that position. Pentameric sequence motifs identified in these aligned recognitions were then clustered by Levenshtein distance [13]. These resulting clusters of unique pentamers and their corresponding pulse duration measurements were evenly distributed across the training, validation, and test sets, ensuring a balanced and representative data partitioning (Fig. 7A).

### SAAV detection with ProteoVue

We developed a comprehensive pipeline for SAAV analysis, called ProteoVue. As illustrated in Figure 8.

**Fig. 8:**
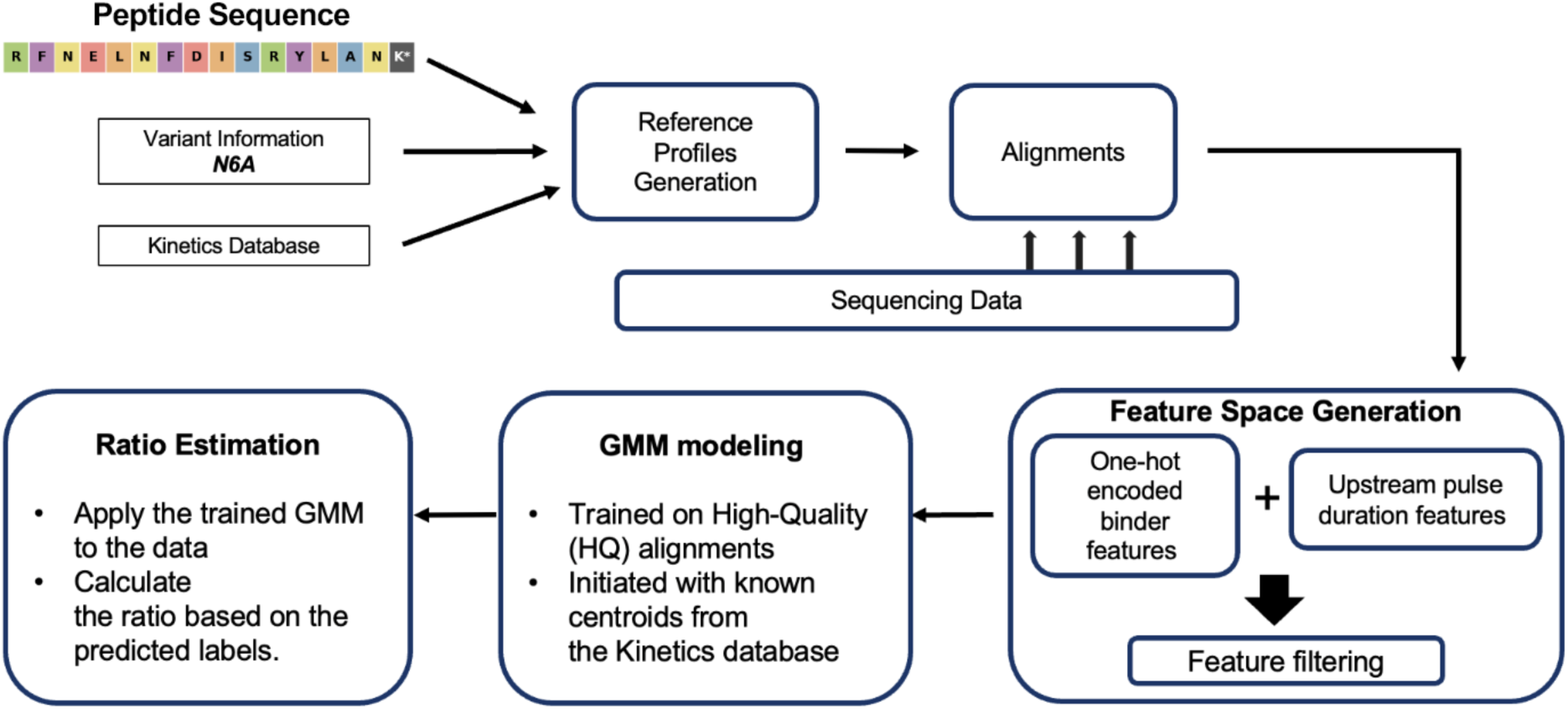
Flow diagram of the steps in ProteoVue.

The workflow begins by generating reference profiles for both peptide variants. Given the peptide sequence, variant position, and substituted residue, ProteoVue retrieves the expected peptide states of the variants and upstream residues, and their corresponding pulse duration profiles from the kinetics database. The reference profile is then used to segment clusters of pulse durations and assign clusters to variants.

Next, sequencing reads from a binary mixture run are aligned to the reference for each of the variants. Only alignments scoring at least 3.75 are used to aggregate positional kinetics, reducing the influence of ambiguous reads. The kinetic features such as PD are then extracted from the aligned reads.

Next, ProteoVue constructs a multidimensional feature space by integrating aggregated positional kinetics—spanning from the variant site up to four residues upstream—with one-hot encoded [14] binary recognizer features. These inputs capture recognizer read variation and the full pentameric context influencing pulse duration. A two-component GMM is trained using these features, with initial centroids guided by the expected kinetic profiles from the kinetic database and recognizer identities.

Applying the trained GMM to the entire dataset yields predicted population identities for all data points. ProteoVue then calculates the ratio of variant populations from the GMM populations.

In summary, ProteoVue accepts raw sequencing data from a binary mixture, a reference sequence, and variant information as inputs, and produces variant population ratio estimates.

### Performance evaluation method

To evaluate the performance of ProteoVue, we calculated the Mean Absolute Error (MAE) between the estimated ratio and the expected ratio across the titrations in the log scale as shown below.

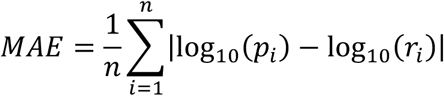

where 𝑛 is the total number of data points, 𝑝_!_ is the predicted ratio of data point 𝑖, and 𝑟_!_is the expected ratio of data point 𝑖. This MAE describes the deviation between the predicted value and the expected value in the log space. A value of 1 in MAE means the estimation ratio is off to the expected ratio by a factor of 10, which is within the expected variability for peptide population predictions given the inherent noise in biological data, model limitations and peptide concentration error. The ability to maintain consistent predictions across a broad dynamic range is more critical than exact agreement at every point.

## Results

### Peptide Alignment - Analysis of Pure Peptides

In this study, we sought to demonstrate SAAV detection in binary mixtures of synthetic peptides. To begin, we sequenced each of the seven variant peptides shown in Figure 2 individually to extract the primary features used by ProteoVue for variant detection. The seven variant peptides were sequenced, and the sequencing data was analyzed using the Peptide Alignment workflow to characterize the kinetic properties of each peptide state in its pure form. The set of reads that aligned to the reference was used to calculate kinetic properties of the various peptide states in that reference, including mean PD, mean IPD, RS start time, and RS duration. Figure 9 displays two example reports for the kinetic properties of the N6 peptide variant (N=177,807 reads). The other six pure peptide samples yield similar alignment reports (see supplementary material). Figure 9B provides additional context by showing the per-read distribution of the various kinetic properties.

**Fig. 9:**
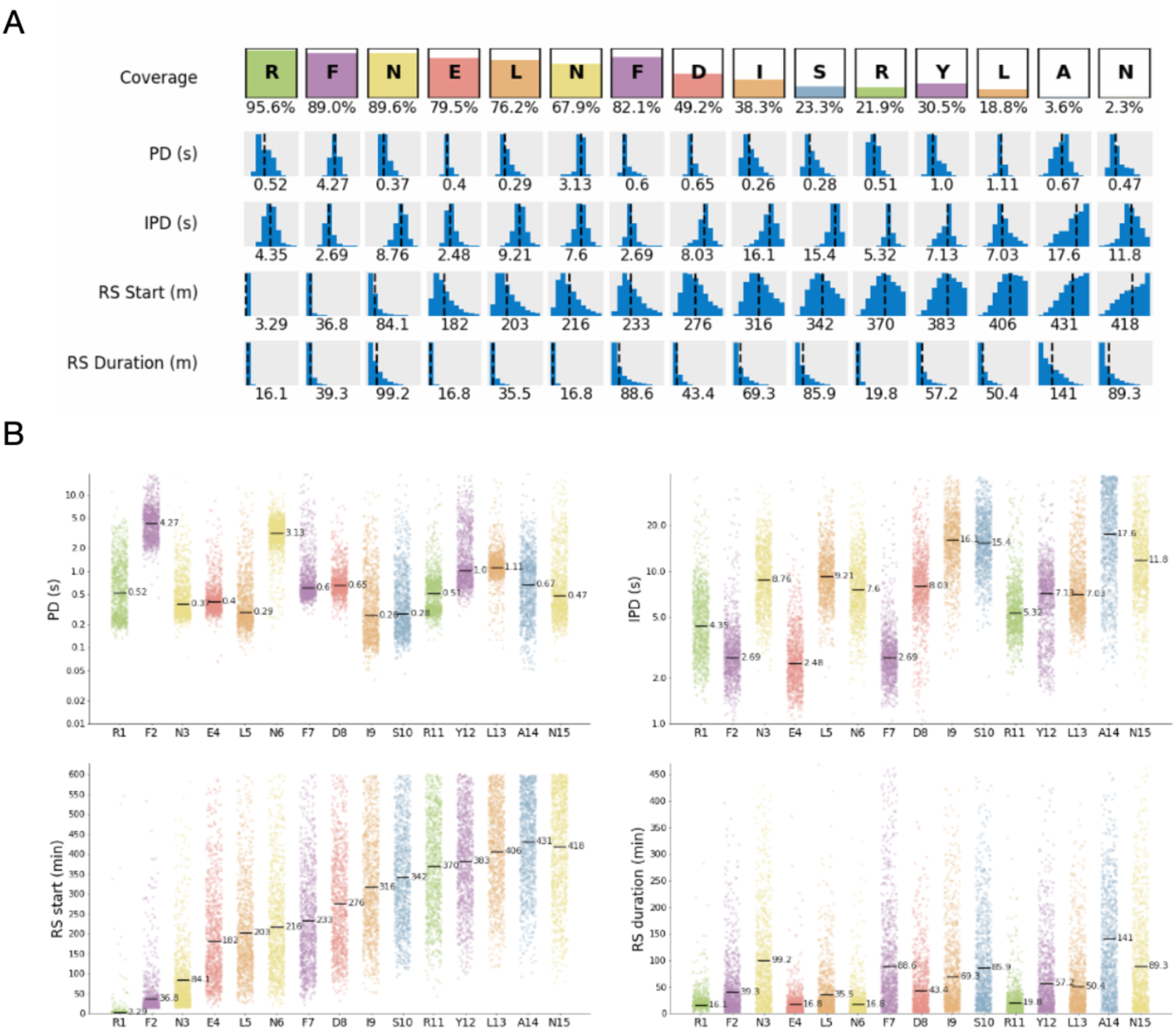
Kinetic properties of the pure N6 peptide. (A) Kinetics summary of aligned reads. (B) Distribution plots of PD, IPD, RS Start and RS Duration.

In addition to these kinetic properties, the Peptide Alignment workflow also provides the coverage for each peptide state. The coverage is the fraction of aligned reads that contain an RS aligned to that state (i.e., the fraction of reads in which that state was not deleted in alignment). Figure 9A displays the coverage of each state. Due to sequencing limitations, the coverage tends to decrease for states deeper into the peptide, with the final two states of the peptide having coverage under 5%.

### ProteoVue

Next we applied ProteoVue to the previously described peptide-mixture titration datasets. Specifically, we analyzed four variants: N to A (N6A), F to W (F6W), R to M (R6M), and C to M (C6M), all occurring at the 6th position of the control peptide (see Materials and Methods for peptide details). For each variant, we analyzed a series of titration datasets with seven distinct mixture ratios—0, 1:100, 1:10, 1:1, 10:1, 100:1, and infinity. The ratios of 0 and infinity correspond to samples containing only one of the two variants. The lowest peptide input was 1nM in 1:100 dilution mixture and was confidently able to detect the variants down to 2pM on-chip. In total, we applied the workflow to 28 samples.

We categorized the variants into three types based on the visibility of amino acids at the variant site. The first type includes visible-to-visible variants (N6A and F6W). The second type includes visible-to-invisible or invisible-to-visible variants (R6M). The third type involves invisible-to- invisible variants (C6M).

In our study, we used pre-selection of primary features for analysis to enhance the accuracy of variant calling. We observed notable differences in the PD at specific upstream positions between the two variant peptides in alignment results from pure peptide samples. We selected the single position that showed the largest difference as the primary feature for each pair of variants. For the variant N6A, we selected position 0, corresponding to the variant site itself, as the primary feature. For F6W, we chose position –4, four residues upstream of the variant site as the primary feature, as it exhibited a twofold difference in pulse duration. For R6M, we chose position -3, three residues upstream of the variant site, as the primary feature due to the most pronounced difference. For C6M, we identified position -1, one residue upstream of the variant site, as the most relevant primary feature.

We evaluated workflow performance by calculating the Spearman’s correlation coefficient (SCC) and the mean absolute error (MAE) between the expected and estimated ratios for each titration dataset.

We first applied the ProteoVue workflow to the N6A titration dataset, using the variant position on the sequence as the primary feature for clustering. Figure 10A illustrates the estimated variant ratios plotted against the expected ratios on a logarithmic scale for N6A. The scatter plot demonstrates a clear diagonal trend, indicative of strong correlation between the estimated and expected ratios (SCC = 96.4%, MAE = 1.198). These observations confirm the detection of variant populations by the workflow.

**Fig 10:**
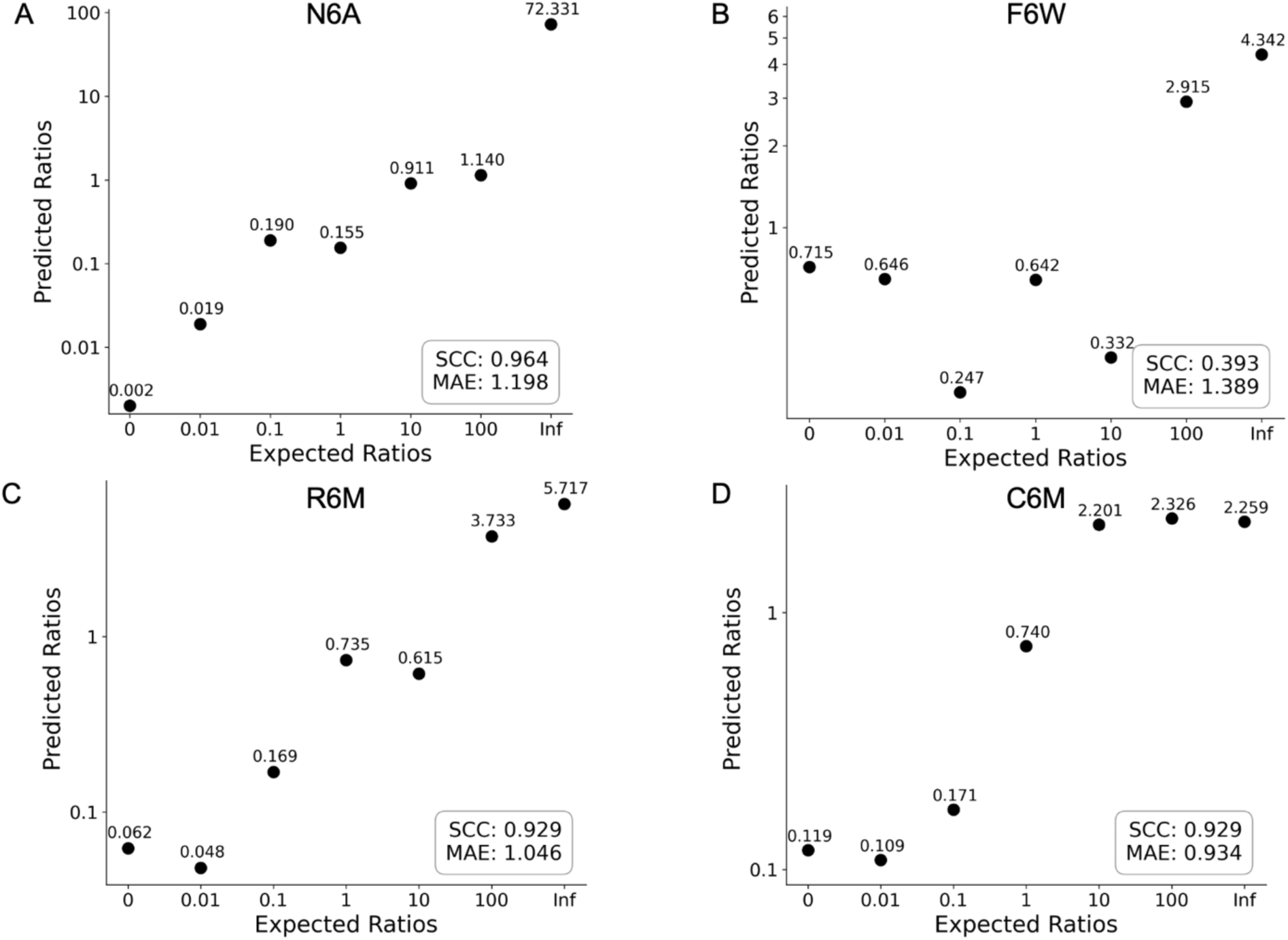
Predicted ratios against expected ratios scatter plot for four variant titration datasets. The x-axis shows the expected ratios between the two variant peptides in log scale and the y-axis shows the predicted ratios in log scale. The value above each dot represents the estimated ratio. The SCC and the log-scale MAE values are shown in the right bottom corner of the plots, indicating the agreement between the estimation and the expectation. Panel A is for variant N6A, panel B is for F6W, panel C is for R6M and panel D is for C6M.

We then applied the same workflow to the remaining three titration datasets—F6W, R6M, and C6M—to evaluate its performance across different types of variants. The estimation-versus- expectation scatter plots are displayed in Figure 10B to Figure 10D.

For the F6W variant (Figure 10B), where both peptides share the same recognizer, the predicted ratios generally align with the expected ratios (SCC=39.3%, MAE=1.389). Although some significant discrepancies occur—due to identical recognizer features limiting alignment and clustering—the features still capture the overall kinetic differences, providing a reasonable approximation of the expected ratios for a challenging variant.

For R6M (Figure 10C), a visible-to-invisible variant, the predictions generally agree with the expected ratios (SCC = 92.9%, MAE = 1.046). Even in the extreme scenario where only the R peptide is present, the slight deviation observed is consistent with individual reads lacking R coverage (see figure S3). This results in a set of reads which are indistinguishable from each other in recognizer features alone, but which can be differentiated by upstream changes in the kinetic features.

The C6M dataset (Figure 10D), representing an invisible-to-invisible variant, posed the most significant challenge. By leveraging the kinetic features, the workflow was able to capture the general trend of the expected ratios (SCC = 92.9%, MAE = 0.934). However, performance deteriorated at extreme ratios, particularly in samples where the expected differences between the two variant peptides exceeded 100-fold. In such cases, the predicted ratios showed minimal differences, likely due to the inherent difficulty in accurately resolving ratios when both peptides are invisible within the system. Nonetheless, the consistent trend observed in the predictions underscores the ability of the workflow to differentiate populations based solely on kinetic features, even in this highly challenging scenario.

We further analyzed the kinetic profiles of the two predicted populations in each pair and across titrations. Figure 11 is an example of the output from ProteoVue showing the kinetic profiles of the reads classified as each variant for N6A at a 1:1 ratio. The pulse duration values exhibited distinct separation not only at the variant site (A6) but also at the upstream L5 and E4 residues, consistent with the expected kinetic effect on upstream positions.

**Fig. 11:**
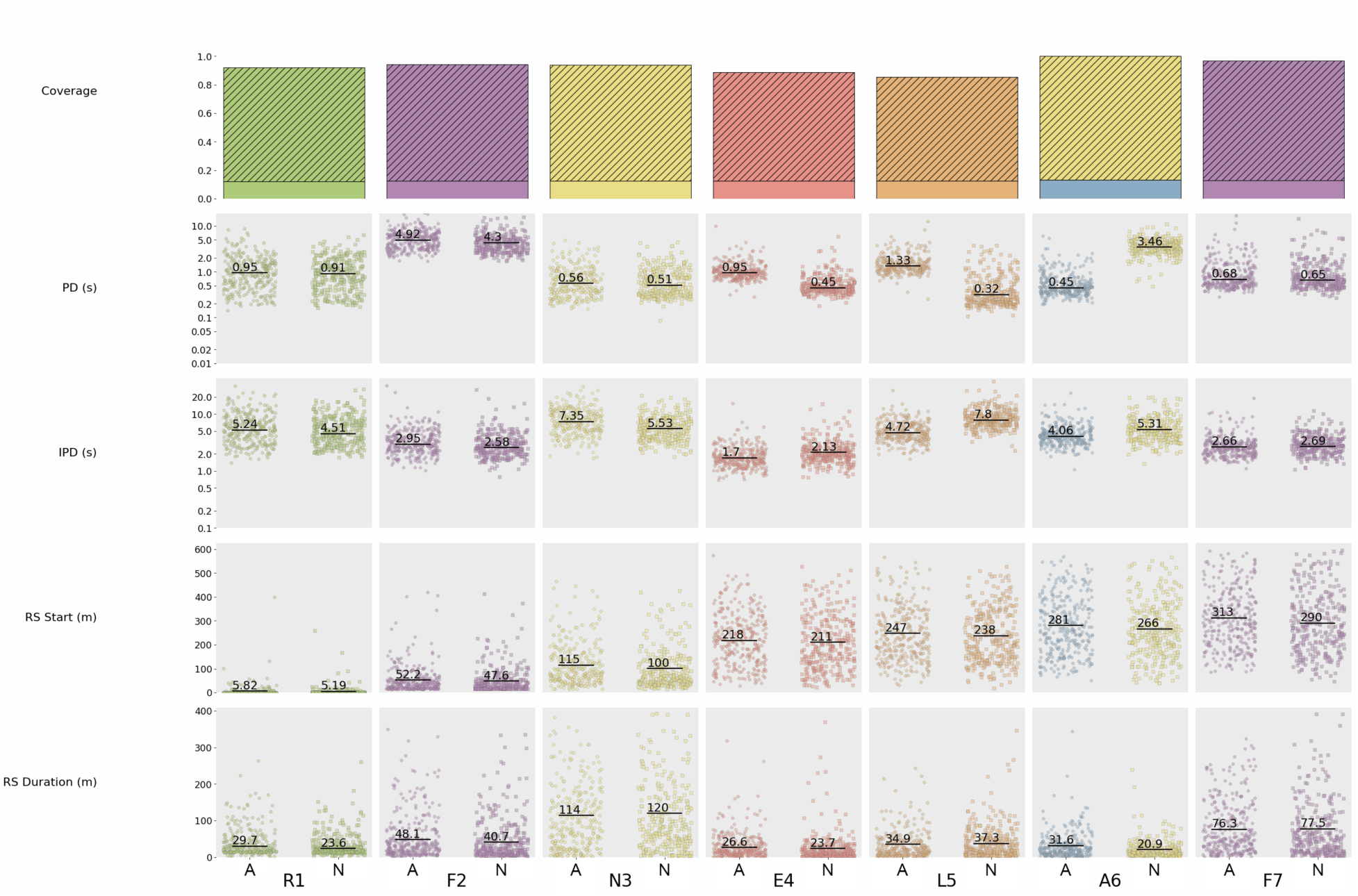
Kinetics profile of the N6A sample at a 1:1 ratio. The A6 variant distributions are on the left and the N6 variant distributions are on the right.

In summary, these results demonstrate that variant calling for SAAVs can be achieved using NGPS. The ProteoVue workflow can estimate the relative abundance of an SAAV to within a factor of 10 while providing distinct kinetic profiles for each population that can be used to further characterize peptide variants. Although we observed specific limitations in extreme cases, the workflow effectively captured the key trends and kinetic distinctions necessary for population differentiation across various types of SAAVs.

## Discussion

In this study, we introduced ProteoVue, a comprehensive bioinformatics pipeline designed for the detection and quantification of peptide variants using NGPS. NGPS leverages single-molecule sequencing to directly measure protein variants at the amino acid level, including SAAVs that are often challenging to detect using conventional methods. By combining advanced signal processing, alignment algorithms, and clustering-based variant calling, ProteoVue addresses key challenges inherent to single-molecule protein sequencing data analysis.

Our results demonstrate that ProteoVue effectively recovers variant peptide populations in controlled mixtures. The output of the workflow correlates well with expected variant ratios across a range of variant types. In particular, the analysis of multiple variants at a defined position revealed that ProteoVue’s kinetic modeling of recognition segmentation can accurately differentiate populations based on subtle differences in single-residue substitutions. Even complex scenarios—such as invisible-to-invisible variants—were resolved, underscoring the pipeline’s utility in extracting information from sparse data.

The ability to handle variants with diverse kinetic and recognizer-binding properties reflects the strength of the underlying alignment and scoring algorithms. The incorporation of a neural network-based kinetic database further refines the scoring, enabling the pipeline to anticipate pulse duration distributions for previously unobserved pentameric sequence motifs. This combination of empirical and predictive modeling ensures that the pipeline remains applicable even as the repertoire of detectable peptide states continues to grow.

Despite these advantages, some limitations remain. Mixtures of variants that share the same recognizer at the variant position pose challenges and sometimes diminish the discriminative power of recognizer-based clustering. Similarly, variants that are fully “invisible” under current conditions can be difficult to distinguish purely based on kinetics, particularly in extreme ratio scenarios. Although the pipeline reliably captures general trends, its accuracy declines when differences between variant populations become pronounced, and both states lack specific recognizer signals. These challenges highlight potential areas for improvement, such as expanding recognizer libraries, refining kinetic models, and incorporating complementary biochemical strategies to increase peptide visibility.

Looking forward, further development of ProteoVue will focus on enhancing the richness and accuracy of its kinetic features. Additional experimental data and improved modeling may better resolve difficult cases such as rare variants or variants in complex proteomes. The integration of more sophisticated statistical frameworks, alternative machine learning methods, and adaptive thresholding could improve variant discrimination in challenging scenarios. Moreover, applying ProteoVue to biological samples rather than synthetic peptides will be a critical step toward evaluating its performance in the context of high sample complexity.

In conclusion, ProteoVue is an important step toward comprehensive SAAV detection and quantification using NGPS. By establishing a pipeline that combines robust signal processing, predictive kinetic modeling, and probabilistic clustering, we have demonstrated the feasibility of reliable variant calling in single-molecule protein sequencing. As NGPS advances, continued development of the bioinformatic approaches in ProteoVue will be essential to unlocking the full potential of this technology in proteomics.

## Author Contributions

Conception and project design: MC and JL

Data acquisition: MC

Data analysis and interpretation of results: JL, EH, MM, DP, KB and IC

Original draft preparation: JL, EH, MM, DP, KB, JV, and IC

All authors reviewed the results and approved the final version of the manuscript.

## Supplementary Information

**Fig. S1:**
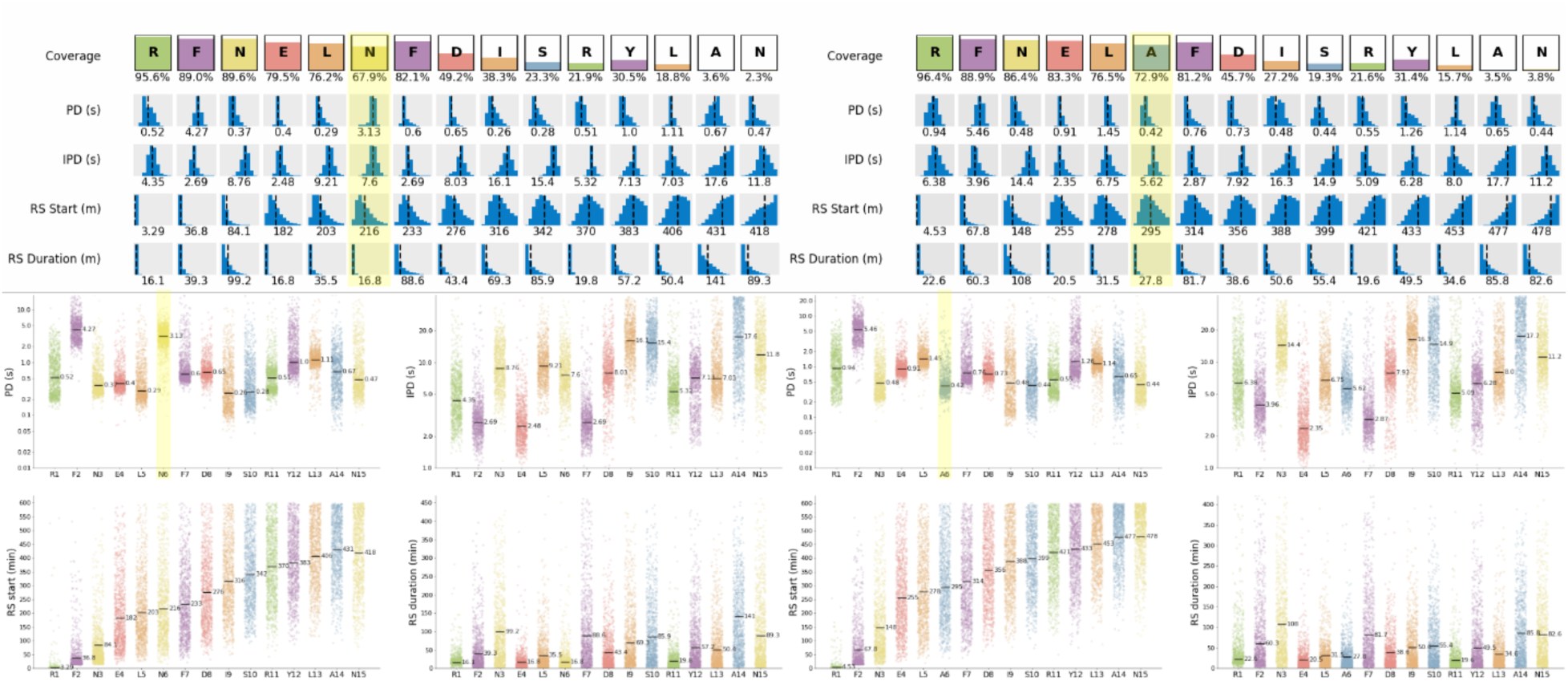
Comparison of the kinetic properties for the N6A variant pair, with the selected kinetic feature highlighted.

**Fig S2:**
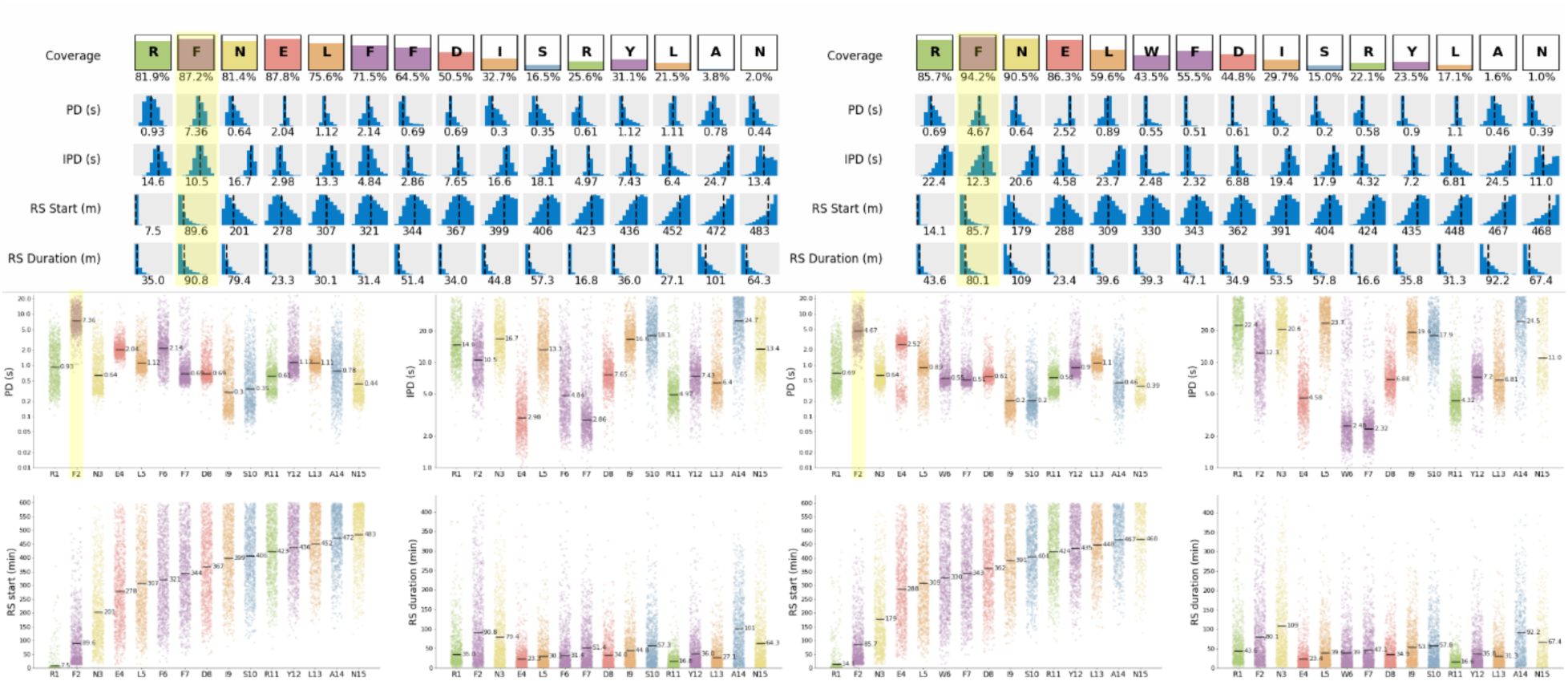
Comparison of the kinetic properties for the F6W variant pair, with the selected kinetic feature highlighted.

**Fig S3:**
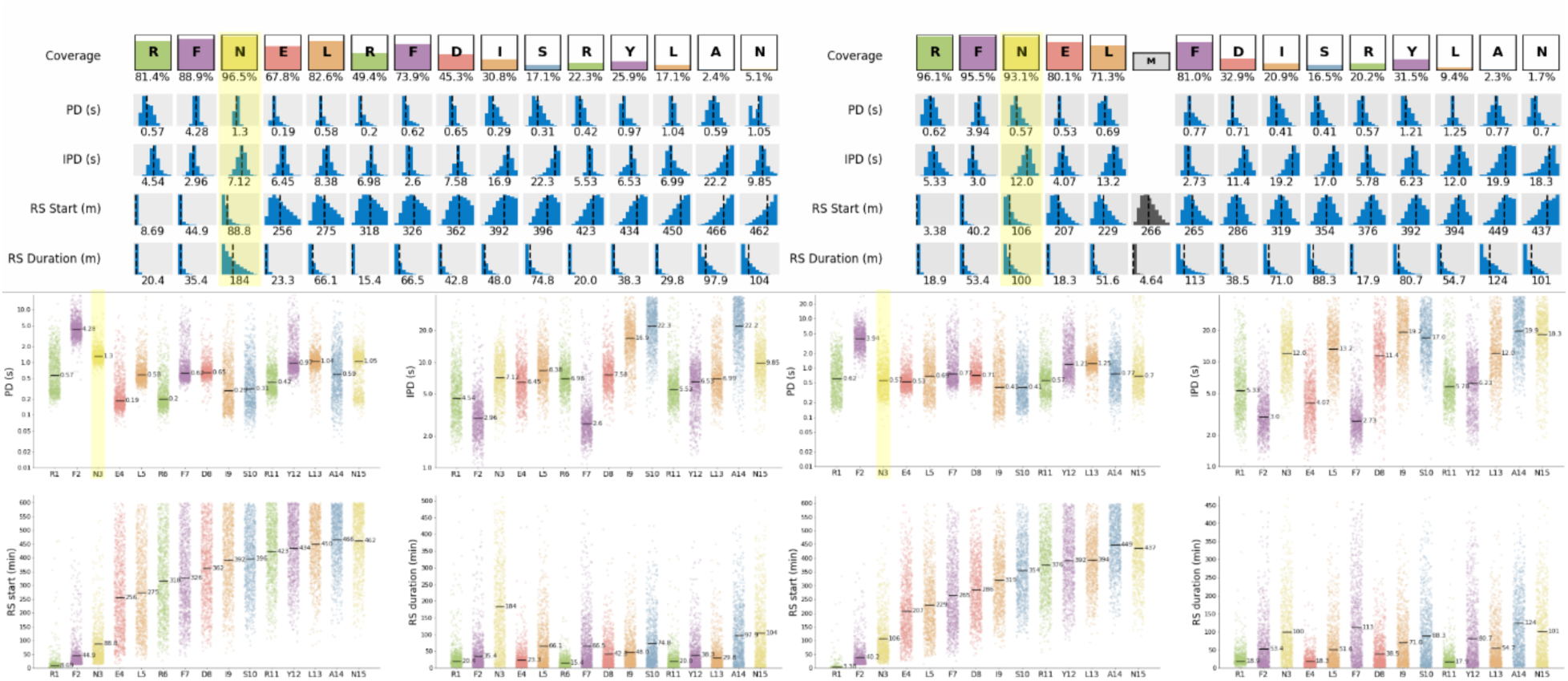
Comparison of the kinetic properties for the R6M variant pair, with the selected kinetic feature highlighted.

**Fig S4:**
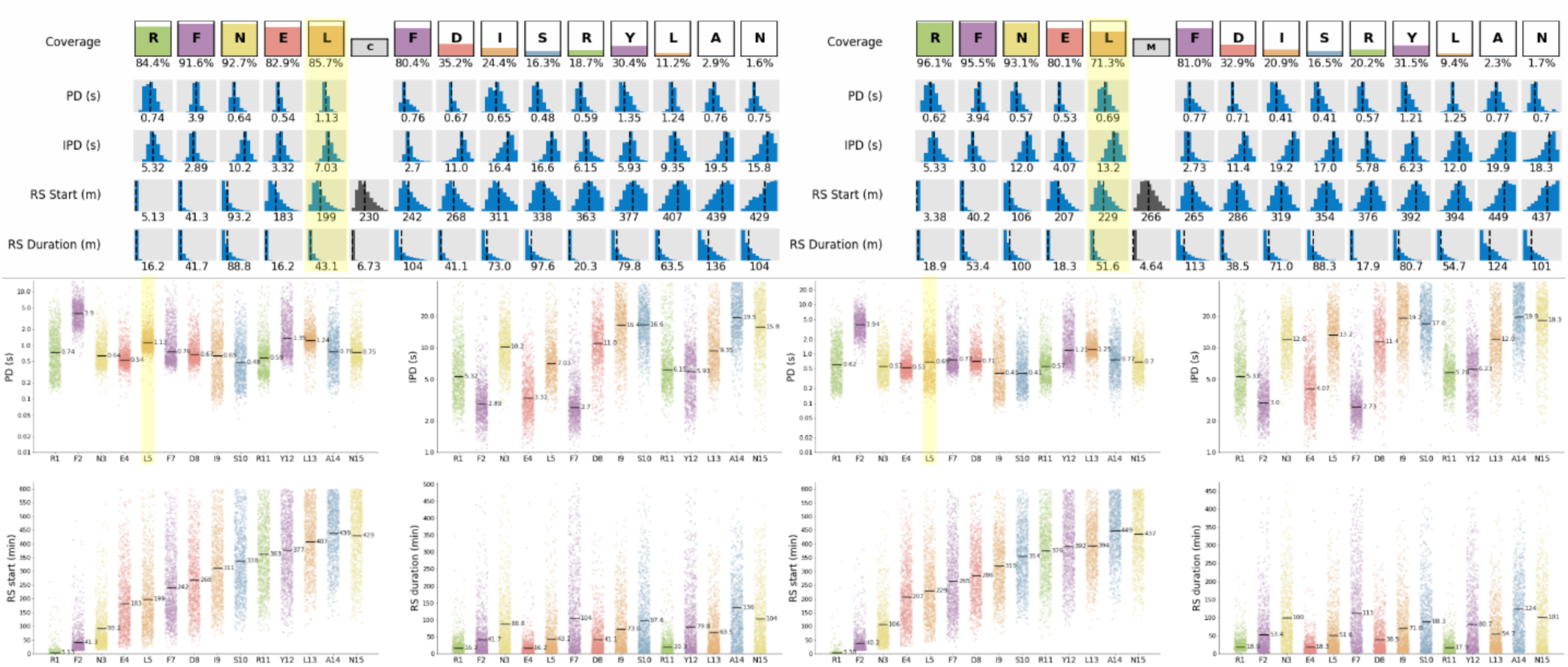
Comparison of the kinetic properties for the C6M variant pair, with the selected kinetic feature highlighted.

